# The timing of natural killer cell response in coronavirus infection: a concise model perspective

**DOI:** 10.1101/2021.08.02.454730

**Authors:** Xiaochan Xu, Kim Sneppen

**Affiliations:** Blegdamsvej 17, 2100, Niels Bohr Institute, Copenhagen University

## Abstract

Coronaviruses, including SARS-CoV, MERS-CoV, and SARS-CoV-2 cause respiratory diseases with remarkably heterogeneous progression. This in part reflects the viral ability to influence the cytokine secretion and thereby the innate immune system. Especially the viral interference of IFN-I signaling and the subsequent deficiency of innate immune response in the early phase have been associated with rapid virus replication and later excessive immune responses. We propose a mathematical framework to analyze IFN-I signaling and its impact on the interaction motif between virus, NK cells and macrophages. The model recapture divergent dynamics of coronavirus infections including the possibility for elevated secretion of IL-6 and IFN-*γ* as a consequence of exacerbated macrophage activation. Dysfunction of NK cells recruitment increase disease severity by leading to a higher viral load peak, the possibility for excessive macrophage activation, and an elevated risk of the cytokine storm. Thus the model predicts that delayed IFN-I signaling could lead to pathogenicity in the latter stage of an infection. Reversely, in case of strong NK recruitment from infected cells we predict a possible chronic disease state with moderate and potentially oscillating virus/cytokine levels.

## INTRODUCTION

Three contagious coronaviruses, severe acute respiratory syndrome-coronavirus (SARS-CoV), Middle-East respiratory syndrome-coronavirus (MERS-CoV), and severe acute respiratory syndrome-coronavirus-2 (SARS-CoV-2), have caused acute lethal diseases in humans. Infection primarily starts in the upper respiratory tract and aggressively progresses to the lower respiratory tract. The patients’ symptoms are heterogeneous from asymptomatic, fever and dry cough to severe pneumonia [1–3]. The severity of illness varies between different groups of people. Among the SARS-CoV-2 patients, the elderly have a relatively higher risk of respiratory failure and case fatality rate [3, 4]. Adults with certain underlying medical conditions (cancer [5, 6], diabetes and obesity [7, 8]) are also at increased risk. In addition, the within-host virus load varies hugely, with 2% of the infected carrying 90% of the virus circulating in the community [9]. It is still under investigation what causes the heterogeneity of the diseases.

Fatal respiratory failure due to lung damage in coronavirus infection is attributed to rapid virus replication and excessive immune responses. The coronaviruses could reach titers as high as 10^8^–10^10^ copies/ml in the upper respiratory tract of patients [10–12] showed by quantification with the pharyngeal swab and sputum samples. Generally, the viral load within patients positively correlates with both the severity of illness and fatal rate [13–15], except in one study with no notable difference between symptomatic and asymptomatic individuals in a young cohort [16]. The spread of the virus is followed by host cell destruction by either virus-induced intracellular stress or by phagocytosis by immune cells contributing to tissue damage [17].

The battle between the virus and the immune system also leads to excessive accumulation of monocyte-derived macrophages in the infected lungs [18], accompanied by highly elevated Interleukin 6 (IL-6) [19]. IL-6 secreted by infected cells starts the accumulation of macrophages by facilitating the recruited monocytes to differentiate into macrophages [20]. The macrophages activated by IFN-*γ* secrete plenty of chemokines and cytokines, including IL-6 and IFN-*γ*, to recruit and activate more macrophages [21]. Thus, macrophages are capable of mediating an excessive potentially self-amplifying innate immune response by positive feedback if activated beyond a tipping point. The over-activated macrophages and the excessive local inflammatory cytokine may cause systemic leukocytes activation. Frequent penetration through the epithelial barrier worsens pulmonary inflammation as substrates from the blood leak into the respiratory tissue. The severe patients suffered from deregulated cytokine production, cytokine storm, and high risk of multiple organ dysfunction syndrome even after the removal of the viruses, which are the main causes of mortality of coronavirus diseases [22, 23].

Innate immunity stops viruses from spreading and prevents the subsequent progression of the disease to later severe stages [24]. While considering the risk of cytokine storm, therapies aiming to exaggerate the patients’ immunity responses should be used carefully. Thus it is vital to understand the dynamics of virus proliferation and its interaction with the host’s innate immune system.

The asymptomatic period of early infections makes it difficult to monitor the infection progression process in the human body, only leaving some reports of PCR counts of mRNA in the upper respiratory tract [16]. Mouse disease models compensate for the paucity of the characterization of early innate immune responses. Strikingly, studies found the dysregulated type I interferon (IFN-I) orchestrates the inflammatory responses in infected mice with SARS-CoV or MERS-CoV [25, 26]. The SARS-CoV-2 also interferes with IFN-I signaling in respiratory cells both in animals and patients [27]. IFN-I is mainly produced by infected epithelial cells first and activated macrophages later. It activates IFN-stimulated genes and antiviral state in bystander cells in a paracrine manner, and some specific immune cells such as natural killer cells (NK cells) [28]. The activated NK cells can recognize and lyse infected cells by their cytotoxicity and activate macrophages by producing IFN-*γ* [29, 30]. Thereby, the interaction between the NK cells and infected cells mediated by IFN-I makes NK cells contribute to the host antiviral state. While the cross-talk mediated by IFN-*γ* between NK cells and macrophages could also be pathogenic by enhancing the excessive activation of macrophages [31]. In clinical observations, NK cells were remarkably recruited and activated in the lungs with SARS-CoV-2 infection [32, 33], which implies their pathogenic role in the macrophage activation as the main producers of IFN-*γ*.

This work discusses the innate immune system as a threshold against early infection events. We build a mathematical model for the interaction network of virus, cytokines, NK cells, and macrophages on the scale of a group of cells. The model reproduces the local excitation of cytokines, and the effect of NK cell function in both virus reduction and activation of macrophages is explored. The model predicts that different levels of macrophage activation could come as a result of heterogeneous NK cell responses. The speed of NK cell responses and subsequent macrophage activation are explicitly elucidated. Our work highlights the importance of proper timing in the treatments of diseases.

## RESULTS

### Dynamical Model of Innate Immune system

The human body has a series of strategies to defend itself against pathogens, broadly classified into innate and adaptive immunity. The fast and direct innate immune responses are particularly vital during early defense against a novel pathogen, while adaptive immunity takes weeks to initiate. In COVID-19 patients, seroconversion appears around 7 days after symptoms onset [12]. As SARS-CoV-2 has an average generation time of 5 [34] to 6.4 days [35], SARS about 8.4 days [36] while MERS has an incubation time of 6 days [37], the adaptive immunity is estimated to establish by two weeks after the coronavirus infection. Comparably, it takes about 12 days before the BNT162b2 mRNA vaccine protects against infections from SARS-CoV-2 [38].

Here we focus on the within-host process of innate immunity combating the virus in the early days after SARS-CoV-2 infection. We simplify the process to a network motif and aim to recapitulate the main features of within-host viral load dynamics, cytokines secretion, and outcomes of disease. NK cells and macrophages (also MΦ) are immune cells considered as representatives of early innate immunity. Generally, the infected cells initiate signaling to trigger immune responses by secreting cytokines including IFN-I, IL-6, and chemokines. Recruited by chemokine gradients, NK cells and monocytes migrates to the infected cells. NK cells are activated by IFN-I directly and induce the apoptosis of infected cells. Thus, they work as sentinels and form a fast counterattack to prevent virus replication. Meanwhile, the recruited monocytes differentiate into pro-inflammatory macrophages mediated by IL-6 with a few days [39]. Further, activated by IFN-*γ*, the macrophages combat the virus by phagocytizing both infected cells and viruses and become the main force against the virus. Both activated NK cells and macrophages could secrete IFN-*γ*. The activated macrophages continue to secrete chemokines and cytokines to recruit other immune cells, including IL-6 to expand the population of themselves (Fig. 1A).

**FIG. 1.**
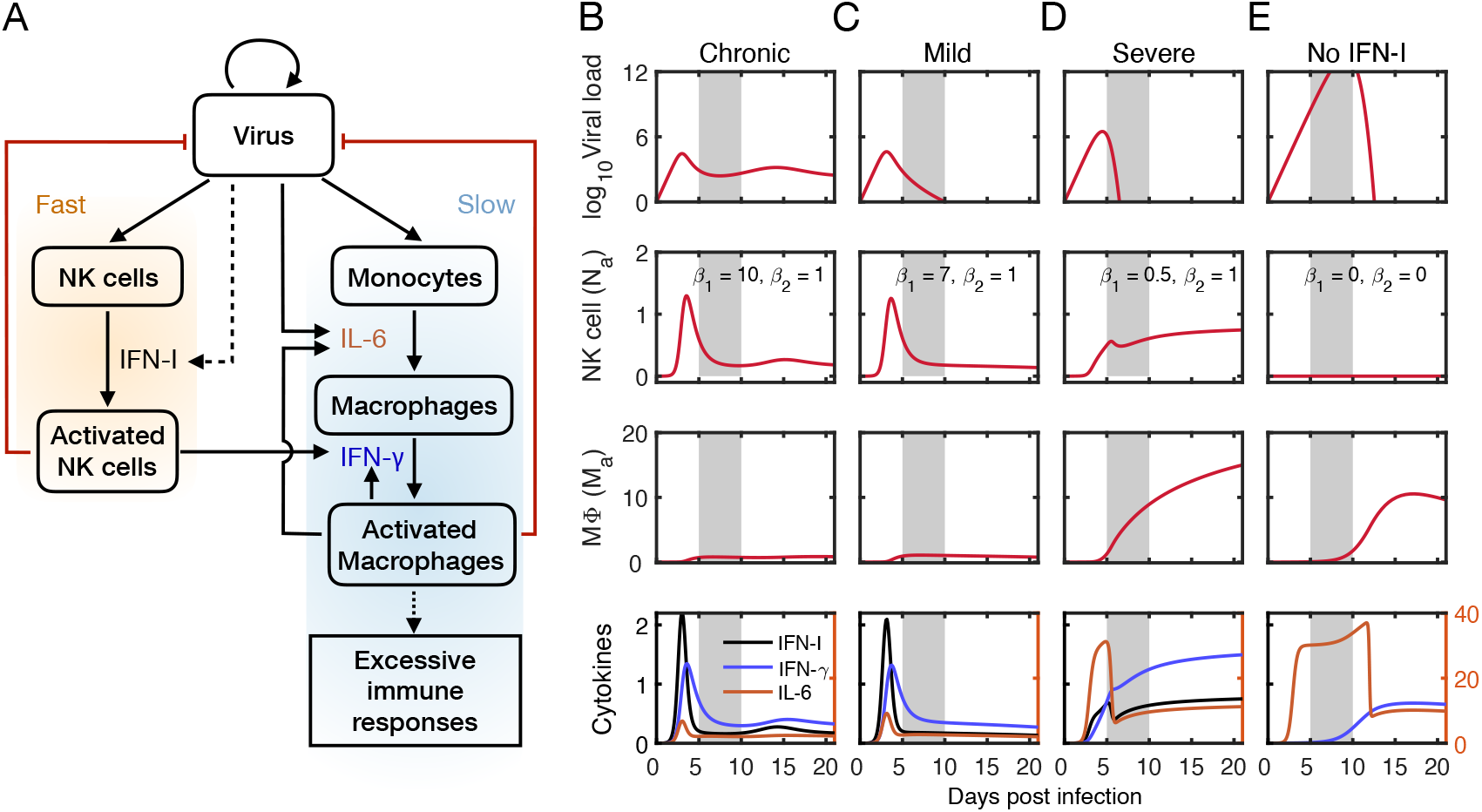
Mathematical model captures outcomes of viral infection at the early stage. (A) Schematic of the immune cells’ combat with the virus-mediated by cytokines. (B–E) IFN-I responses to viral infection affect the outcomes of illness and lead to different cytokines secretion features. (B) Chronic viral load dynamic with prompt IFN-I signaling response (*β*_1_ = 10, *β*_2_ = 1). (C) Mild symptom with normal IFN-I signaling response (*β*_1_ = 7, *β*_2_ = 1). (D) Severe illness with insensitive IFN-I signaling response (*β*_1_ = 0.5, *β*_2_ = 1). (E) Severity of disease reduced by early treatment with IFN-I blockade (*β*_1_ = 0, *β*_2_ = 0). Other parameters in the model: *K*_1_ = 10^5^, *K*_2_ = 5, *γ*_1_ = 30, and *γ*_2_ = 15. Initial viral load is *V*_0_ = 1, other variables initiate at 0.

A moderately detailed model with four variables, the virus load (*V*) also serves proxy for the number of infected cells, the activated NK cells (*N_a_*), the monocyte-derived macrophages (*M_m_*), and the activated macrophages (*M_a_*) describes the above processes. Considering a small region of the body, for example, a small group of the epithelial cells in the upper respiratory tract, the model follows the progress of this local infection by exposure to an initial virus load *V*_0_. The virus grows exponentially with a replication rate 1/τ*_C_*, unless counteracted by activated NK cells (*N_a_*) and macrophages (*M_a_*). The virus replication time scale within pulmonary cells is estimated to about 6 hours [40], τ*_C_* = 6, in the equations. Apart from an earlier onset of disease, the results are robust to changing this parameter to 4 hours, possibly reflecting a higher *in vivo* spreading rate of the virus. The half-life of cytokines is as short as minutes [41]. Thus, we assume instant adjustment of cytokines to the current level of secreting cells:

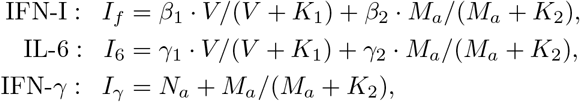

where the last equation expresses that IFN-*γ* mainly comes from NK cells and activated macrophages. *K*_1_ and *K*_2_ set the thresholds for maximal activation from virus and macrophages, and the terms *V*/(*V* + *K*_1_) and *M_a_*/(*M_a_* + *K*_2_) reflect that secretion of cytokines is limited. For simplicity, we use the same threshold for activation of the different cytokines. We did not include rate parameters for INF-*γ* as these can be absorbed in the other rate parameters.

In other infections, NK cells are rapidly and directly recruited to the infected site within 24 h and peak by 3 days after infection [42, 43]. We use a similar time scale with τ*_N_* = 24 h for the dynamics of activated NK cells here. Unlike the recruited NK cells, the monocyte-derived macrophages go through a few steps before becoming efficient defenders. The accumulation rate of macrophages here is mainly affected by the differentiation from recruited monocytes to proinflammatory macrophages with a relatively slow time scale indicated as τ*_M_* = 100 h.

The model is

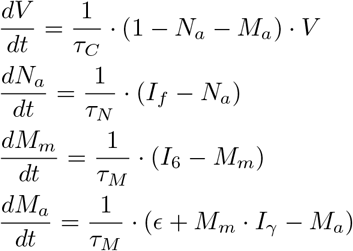

where we measure the level of *N_a_* and *M_a_* in units of their respective ability to kill the virus and infected cells.

In reality, macrophages can also be activated through other paths [44], quantified by a relatively small constant *ϵ* = 0.05 in our simulations. We simulate the process for 20 days after the viral infection and focus on the early dynamics before adaptive immunity.

### Severity of disease dependent on the timing of NK cell activation

The blocked IFN-I signaling is a common feature in the three coronavirus diseases, which may explain the overall success of the viral invasion. We first explored what happened if the activation of NK cells is affected with modulation of IFN-I signaling. In our model, the parameter *β*_1_ is representative of the modulation. Larger *β*_1_ indicates the high sensitivity of the NK cells to virus invasion, while smaller *β*_1_ indicates the IFN-I signaling is inhibited and failed to activate the NK cells.

The model predicts different outcomes of infection with various sensitivity of the NK cell activation to viral stimuli (Fig. 1B-E). On the whole, high sensitivity leads to prompt NK cell response initiated by the immediate secretion of IFN-I, preventing the virus from reaching high numbers (*β*_1_ = 10 and *β*_1_ = 7, Fig. 1B-C). Low sensitivity leads to delayed NK cell response, higher viral load peaks, and elevated macrophage activation levels (*β*_1_ = 0.5, Fig. 1D). The different dynamics of the virus and immune cells mimic different severity of the disease in days 5–10 after infection. The infected host has light symptoms and a low activation level of macrophages (Fig. 1B-C), corresponding to asymptomatic and mild clinical observations. All the cytokines and NK cells increase with the viral replication and decay with the viral clearance by the immune cells. This spike-like pattern indicates the virus is dominant in driving the inflammation in these situations. Interestingly, we found the prompt NK cell response can introduce a persistent within-host viral dynamic which bears the similarity to some cases where SARS-CoV-2 fails to cause strong symptoms but anyway persists in the host [45]. We will discuss the chronic pattern later.

The severe infected host has severe symptoms with continually increasing macrophage activation even after the viral clearance (Fig. 1D) accompanied with the constant secretion of cytokines. IL-6 rather than IFN-I becomes the first elevated cytokine and keeps at a high level. The NK cells are recruited and activated by macrophages at the late period. These dynamics indicate that the macrophages drive the excessive inflammation after the virus triggers immune responses. The high viral load may increase the spread from the local site, which may cause wilder scale damage through direct destruction of tissues and intensified inflammation.

In mice models with SARS-CoV or MERS-CoV infection, blocking IFN-I signaling decreased macrophage accumulation and protected the animals [25, 26]. To reproduce the experiments of abrogation of IFN-I signaling, we block the NK cells by setting *β*_1_ = 0 and *β*_2_ = 0. The macrophage activation is lower than the severe scenario and decreases naturally even if the viral peak achieves higher (Fig. 1E). In the model, it is due to reduction of the macrophage self-amplification effect by inhibition of NK cell function, which explains the experimental result that infected mice are protected by blocking IFN-I signaling from fatal illness.

The prediction that different activation timing of NK cells leads to different outcome of diseases is suitable for discussion in many contexts. Except that different viruses have different ability to interfere IFN-I signaling, it also explains various immune response strength in the hosts. In the same person with the infection of a given virus, different tissues may have different IFN-I responses, leading to heterogeneous spread. In diabetic patients, different degrees of the impaired NK cell function [46] correspond to different *β*_1_. Thereby, the parameter *β*_1_ could quantify the possible differences between patients besides variations between infection sites inside a person. Immunodeficient patients may have fewer NK-cells, which may also explain a worse prognosis for these types of patients [47]. Overall, our model suggest various venues for explaining the heterogeneity of disease, all associated to a change in the early recruitment or in the overall/local capacity of relevant immune cells.

### Protective and pathogenic roles of NK cells in the disease progression

From the basic model, we found that proper timing of NK cell response is important to attenuate the risk of excessive accumulation and activation of macrophages. It may be further analyzed by approximating *M_m_* ≈ *I*_6_:

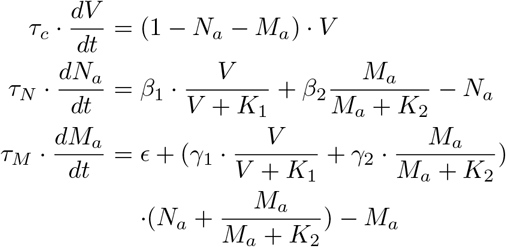

Importantly, this simple model maintains the hierarchy in timescales, τ*_C_* ≪ τ*_N_* ≪ τ*_M_* (Fig. 2A). This model with three variables exhibits similar infection trajectories as the more detailed model. It further illustrates that the higher initial viral load leads to faster disease progression and slightly more severe disease (pink versus the red trajectories in Fig. 2B–D).

**FIG. 2.**
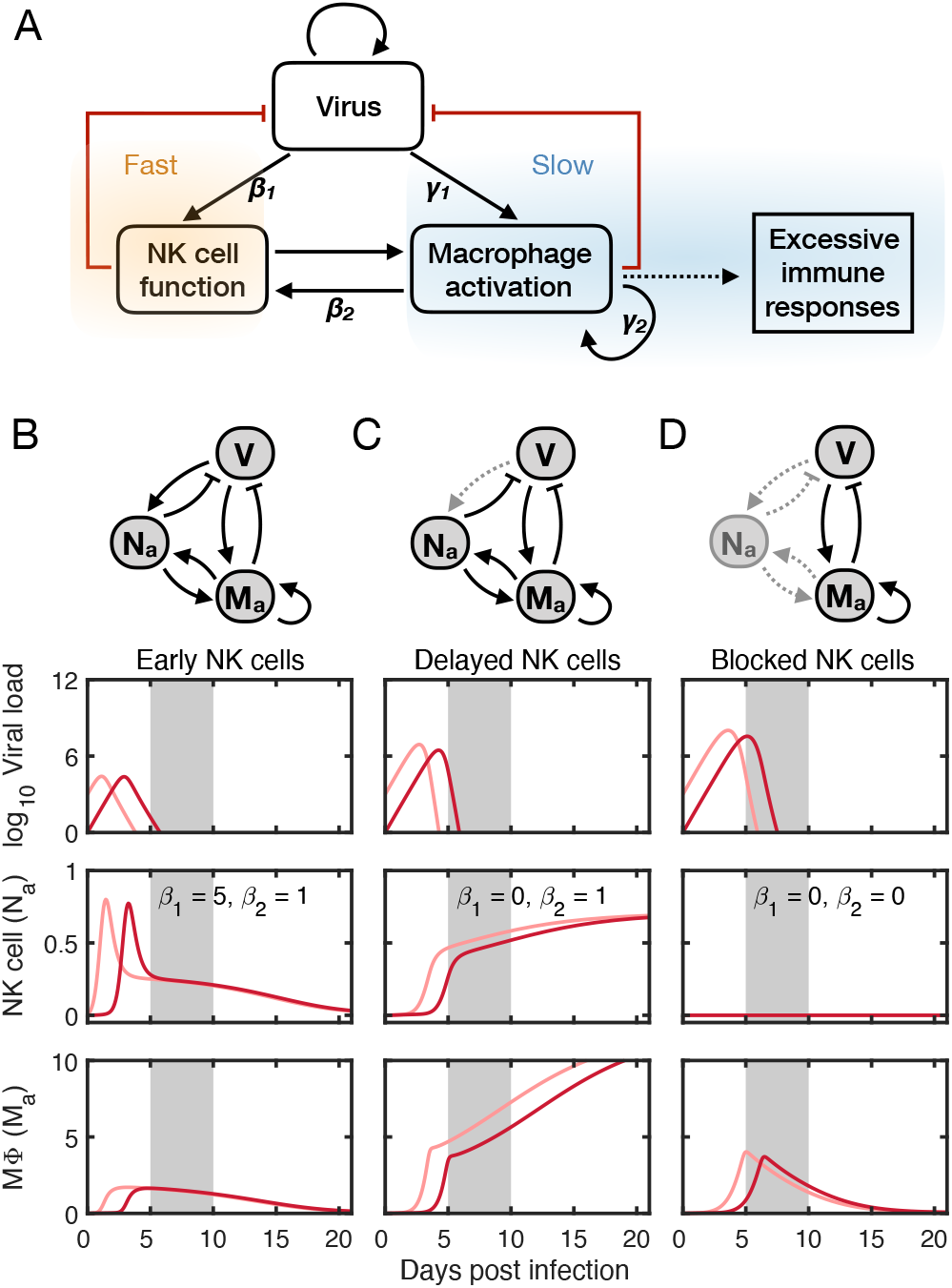
Simplified mathematical model captures the dual roles of NK cells in virus-triggered macrophage activation. (A) Schematic of virus, NK cell function, and macrophage activation. (B–D) Dynamics and outcomes of different NK cell responses to viral infection. (B) Mild macrophage activation with early NK cell response (*β*_1_ = 5, *β*_2_ = 1). (C) Exacerbated macrophage activation with delayed NK cell response (*β*_1_ = 0, *β*_2_ = 1). (D) Reduced macrophage activation by blocking NK cells (*β*_1_ = 0, *β*_2_ = 0). Other parameters in the model: *K*_1_ = 10^5^, *K*_2_ = 5, *γ*_1_ = 20, and *γ*_2_ = 12. Initial conditions: Red, [*V, N_a_, M_a_*]_*t*=0_ = [1, 0, 0]; pink, [*V*, *N_a_*, *M_a_*]_*t*=0_ = [10^3^, 0, 0].

After the virus clearance, the NK cell response follows macrophage dynamics due to a much slower time scale of macrophages. After virus clearance (*V* = 0) the model could be further simplified by using the fact that NK cells are fast to adjust, *N_a_* = β_2_ · *M_a_*/(*M_a_* + *K*_2_):

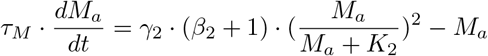

This has the steady state solutions:

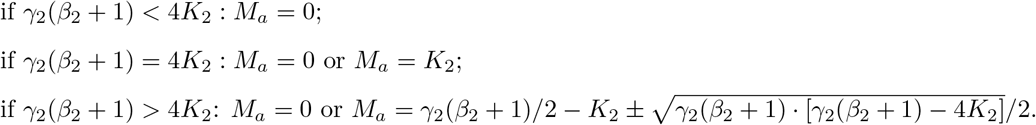

Thus, the motif for the interaction of NK cell and macrophage has a saddle-node bifurcation at *β*_2_ = 4*K*_2_/*γ*_2_ − 1, see Fig. 3A). With small *β*_2_, macrophages are temporally activated then decay to *M_a_* = 0. With large *β*_2_, besides decay to 0, macrophages could establish self-amplification, and the host probably suffers from a cytokine storm. Thus the parameter *β*_2_ captured the pathogenic role of NK in the macrophage activation-induced cytokine storm.

**FIG. 3.**
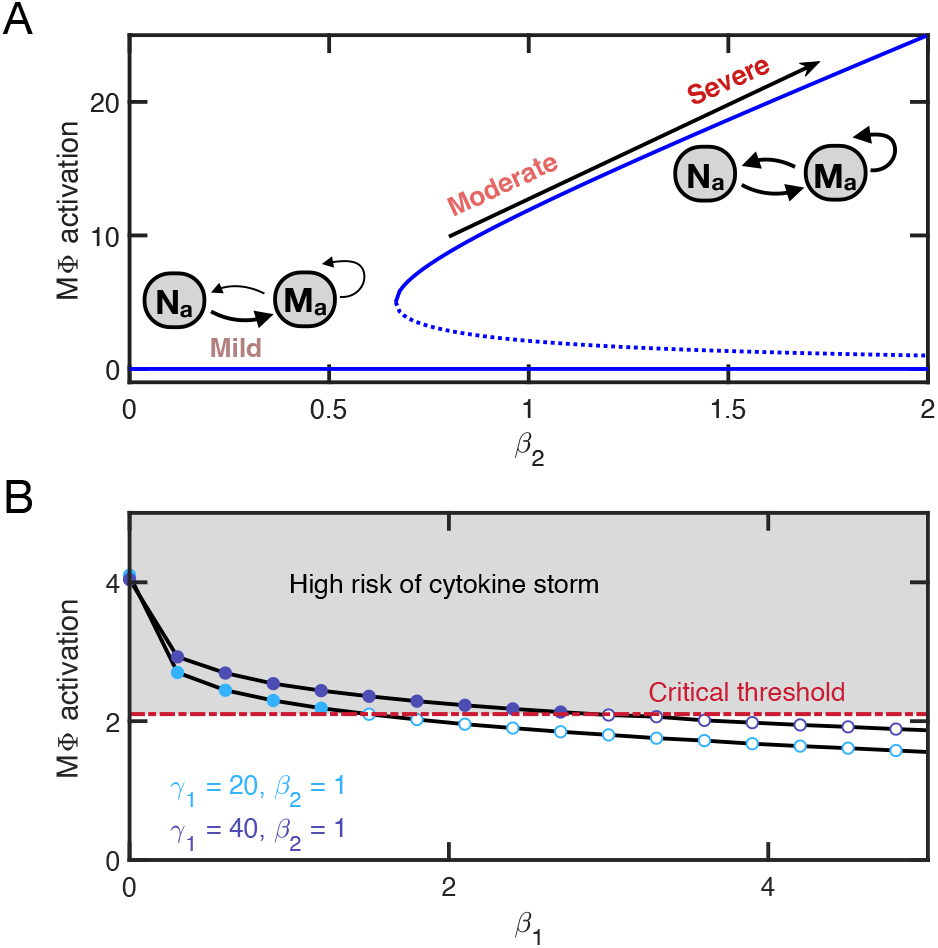
Severity of disease dependent on protective and pathogenic effects of NK cells. (A) Pathogenic effect of NK cell characterized by *β*_2_, increased *β*_2_ can trigger a saddle-node bifurcation and lead to higher disease severity. Stable (solid) and unstable (dashed) steady states of macrophage activation are calculated with corresponding *β*_2_. (B) Protective effect of NK cells characterized by *β*_1_, increased *β*_1_ can prevent the macrophage from being over-activated and leading to cytokine storm. The macrophage activation level at the time when the invading virus is removed (y-axis) is compared with the critical threshold. Filled, activation levels putting the host at high risk for developing cytokine storm; hollow, activation levels indicating the host’s recovery later on. Blue, *γ*_1_ = 20 and purple, *γ*_1_ = 40.

With large *β*_2_, the macrophage activation level at *V* = 0 resulted from interactions between virus, NK cells, and macrophages in the early battle decides if the host recovers or has a high risk of a cytokine storm. The host recovers from the infection only if the macrophage activation achieves a level below a critical threshold when finishing the virus clearance. Otherwise, the host is at high risk for a cytokine storm mediated by activated macrophages.

The critical threshold corresponds to the unstable steady state with the given *β*_2_, *M_a_* = γ_2_(*β*_2_ +1)/2 − *K*_2_ − 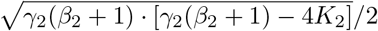. The parameters *β*_1_ and *γ*_1_ become determinant in the early phase. The larger *β*_1_ is, the lower level of macrophage activation at *V* = 0 (Fig. 3B), indicating the defensive role of NK cells in the disease progression. Meanwhile, the macrophage activation also increases with the parameter *γ*_1_ quantifying how sensitive the macrophages’ accumulation is to viral replication through IL-6 (Fig. 3B).

### Chronic pattern and recurrence of disease caused by strong NK cell function

A small proportion of the patients with coronavirus infection suffered from persistent symptoms (long-term COVID-19). The long-term COVID-19 is mainly caused by tissue function impairment as complications of COVID-19 at the acute disease phase [48]. In some cases, the patients were asymptomatic or had mild symptoms first but later found to carry persistent and recurrent virus [49, 50] before the onset of severe disease. We thus explored in which scenario the interaction between the virus and immune system leads to a chronic pattern with persistent or recurrent viral dynamics but without severe symptoms.

We found the chronic pattern could be caused by NK cells with a strong reaction to the virus that directly reduces the viral load. In this case, a short-term stimulation from the virus prevents the macrophages from accumulating. The infected person will be asymptomatic and show a persistent viral load (Fig. 1B) or re-positive clinical feature (Fig. 4A), which are disease progressions observed.

**FIG. 4.**
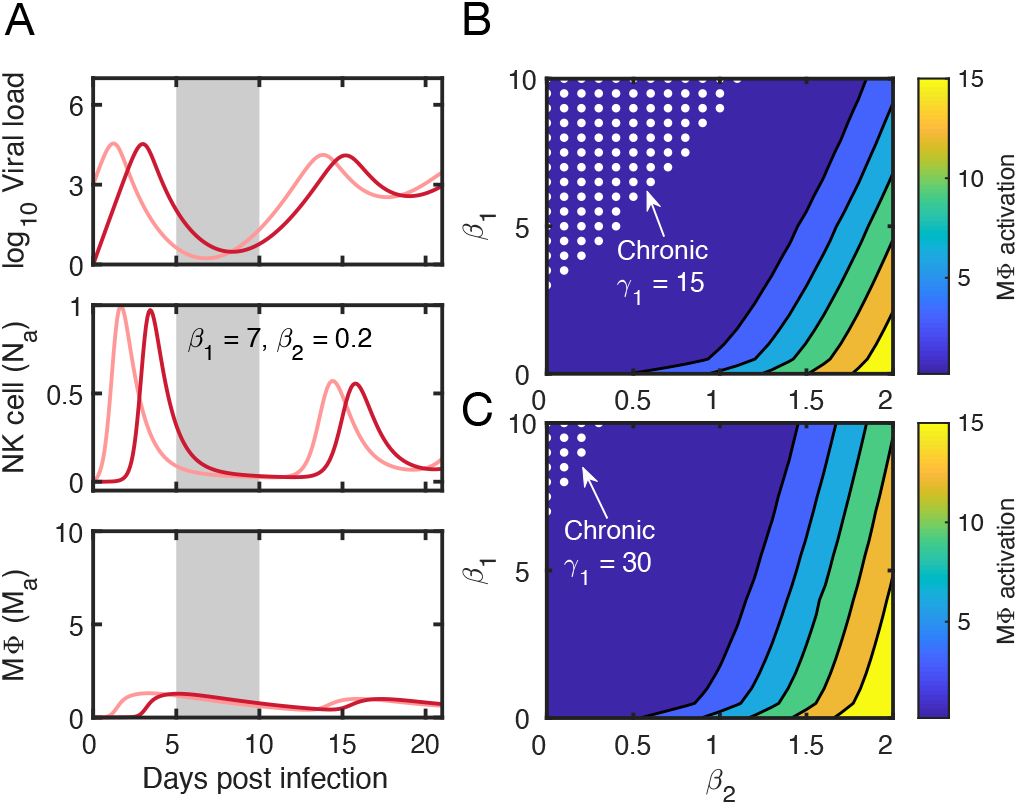
Persistent low viral dynamics with strong NK cell function. (A) Dynamics of recurrent virus and reactivated NK cells. Initial conditions: Red, [*V, N_a_, M_a_*]_*t*=0_ = [1, 0, 0]; pink, [*V*, *N_a_*, *M_a_*]_*t*=0_ = [10^3^, 0, 0]. (B-C) The range of protective function (*β*_1_) and pathogenic function (*β*_2_) of NK cells which causes chronic pattern of the disease. The white dots indicates the parameter pairs of *β*_1_ and *β*_2_ with *γ*_1_ = 15 (B) and *γ*_1_ = 30 (C) respectively. Color key: macrophage activation level at Day 20 indicating the severity of symptoms and risk for cytokine storm.

Furthermore, the parameter *γ*_1_ is a determining factor for chronic development, reflecting the importance of macrophages’ sensitivity to the virus. The parameters range of *β*_1_ and *β*_2_ causing chronic pattern of the within-host viral dynamics shrinks when *γ*_1_ is larger (Fig. 4B and C).

The model predicts that chronic disease development occurs when *β*_2_ is small while *β*_1_ is large. It indicates the cytotoxicity of NK cells plays a dominant role in virus clearance as the activity of the macrophages is limited. These results also suggest that for patients with mild symptoms, using IFN-I treatment alone may not be enough to thoroughly remove the virus, especially in elderly patients with a decrease in macrophage function [51]. While applying IFN-I in combination with other therapeutic strategies, including those with immunosuppression effects, one should also consider the delicate balance between NK cell function and macrophage activation.

## DISCUSSION

Coronavirus infection activates virus-and-host interactions that vary between persons and between tissues within persons. This paper introduces a framework to the earliest manifestations of these attack and defense mechanisms: The innate immune system of NK cells and macrophages recruited to a relatively localized infection site. Any successful pathogen will likely have ways to counteract this recruitment, with coronavirus reported to invoke mechanisms associated to manipulation of cellular stress and a reduction of the direct NK cells recruitment from infected cells.

Our model highlights the implications of the dysfunction of NK cells in the innate immunity caused by the insensitivity of IFN-I response to viral infection. Meanwhile, any immune deficiency converged to NK cell functions could be discussed with our model. Our model can also be applied in some other contexts when a transient player appears and remits the pressure of the immune system caused by antigens. The idea of multiple layer effectors and different time scales of immune responses are also inspiring for similar situations. This part of the model would be valid for a larger class of infecting viruses, and in general, predicts increased vulnerability for the elderly. In contrast, children have evaded COVID-19 [52] and consistently had weaker symptoms, perhaps reflecting a general tendency of having a faster reacting innate immune system [53]. The NK cells already migrate to the infected tissues by a hint of detected decrease in circulation in the early acute disease phase in infected children with mild symptoms, which does not happen in adults[54]. It indicates the NK cells respond more promptly in children when the virus invades, corresponding to larger *β*_1_ in our model. The model partially explained the unaccountable observation of the difference between children and adults. The higher sensitivity to the viral infection by the NK cells would limit the replication of the virus in the early acute disease phase.

In reality, the host is exposed to widely different viral loads at infection, dependent on the situation. A high initial load may lead to a worse prognosis. In our model, we found the maximum virus load positively correlates with the macrophage activation. The outcome is more severe with an increased initial viral load, achieving a slightly higher viral peak load and slightly faster macrophage activation. It means that the infected person may become more infectious. The overall faster progression of the disease means that the adaptive immune response comes relatively later in the disease progression, allowing for more damage.

Our model has limitations and should only be seen as a modest step of putting key elements of the innate immune system into a dynamic framework. In particular, we omit the anti-inflammatory cytokines secreted by the macrophages that reduce recruitment of macrophages in a previous model of innate immunity [55]. Extension of these next-level cells may reduce and extend the macrophage dynamics to the longer timescales where adaptive immunity also comes into play. The model could be extended further into the disease progression with more components in the immune systems including the role of CD8 T cells. A previous study about the motif comprised of the interactions between antigen and CD8 T cells also suggested that bistability would result from the interplay between antigen and the immune system [56].

It is also interesting to compare our model to the quite different and elaborate model of ref. [57] that follows a population of human cells by assuming that IFN-1 converts infected cells to resistant cells. In that model, the NK cells are the main cause of direct virus elimination, and the macrophages primarily act by inducing IFN-1 that in turn reduce virus proliferation. Despite the substantial differences, the model in ref. [57] contains the central motif of Fig. 2B, leading to similar predictions of increased macrophages activity associated with reduction of IFN-1 signaling.

The heterogeneity of the viral kinetics has been discussed in other relevant models in contexts that included contact networks [58] and COVID-19 treatments [59, 60]. Predicted by these models, the virus population peak around symptom onset and its subsequent decay are the main elements of heterogeneous disease kinetics. A larger and detailed full-scale model [61] simulates the viral infection dynamics at the whole-body level while highlighting the crucial roles of innate immunity. In this light, the contribution of our model is its simplicity and focusing on the early blockade to the virus replication by the innate immunity. Perhaps the most important comparison is to consider disease trajectories in one study [16] indicating that the virus load often increase by about a factor 10^5^ in 3 days before its peak value, and thereafter it takes about 6 days for viral mRNA to decrease by a similar factor. The observed decline is in discord with our typical prediction of a rapid decrease of virus count for the more severe cases (Fig. 1D, E and Fig. 2C, D). This discrepancy may in part reflect that our model only considers a single infection site, which may differ substantially from the more global progression between body parts. The virus peak and its subsequent decline should in principle be augmented by a spatial model that takes into account infections in the separated upper and lower respiratory tract, thus leading to a longer disease trajectory than we typically predict.

Previous models focus on the period around and after the peak of virus shedding. This makes the models easier to calibrate against available clinical data that is almost only available after symptoms onset. Data for the virus/cytokine kinetics before symptom onset is still lacking. Our model suggests a way to organise our thinking about the intricate and presumably noisy early infection dynamics. We predict that heterogeneity appears and is enlarged before symptom onset. As demonstrated, the multiple levels of attack and defence mechanisms provide multiple venues for heterogeneity in disease development, including pronounced heterogeneity in both virus shedding and subsequent symptom severity.

Our model gives a general way to think about how to alter treatment of the patients who are at different stages when admitted to hospital. The protective and pathogenic roles of NK cells need to be delicately balanced when targeting IFN-I in combination with other therapies.

## ACKNOWLEDGMENTS

We thank Ala Trusina for enlightening discussions. We also thank Mikkel S. Svenningsen and Julius B. Kirkegaard for constructive comments. This project has received funding from the European Research Council (ERC) under the European Union’s Horizon 2020 research and innovation program under grant agreement No 740704.

## AUTHOR CONTRIBUTIONS

X.X. and K.S. designed the study, performed the research and wrote the manuscript. X.X. implemented models and all the simulations. All authors reviewed the manuscript.

## DATA AVAILABILITY

The datasets generated during and/or analysed during the current study are presented in the manuscript.

